# A Modular Chemoenzymatic Approach to C14-Functionalized Steroids

**DOI:** 10.1101/2022.06.08.495276

**Authors:** Fuzhen Song, Mengmeng Zheng, Junlin Wang, Huanhuan Liu, Zhi Lin, Benben Liu, Zixin Deng, Qianghui Zhou, Xudong Qu

## Abstract

C14-functionalized steroids belong to a unique class of steroids with important biological activities. However, the lack of efficient methods to access C14-functionalized steroids impede related steroidal drug discovery. Herein we report a modular chemoenzymatic approach to access diversified C14-functionalized steroids. We first identified a novel C14α-hydroxylase (CYP14A) from *Cochliobolus lunatus* with high catalytic efficiency and substrate promiscuity. Protein engineering of CYP14A generated three variants I111A, M115K and V124A that greatly improved the C14-hydroxy regioselectivity. Based on this efficient biocatalytic method, a range of C14α-OH steroids with C17 side chain were prepared in good yields, which was then transformed into Δ14 olefins through a facile elimination. The newly formed Δ14 olefin served as a versatile handle to install diversified functional groups (e.g. epoxide, β-OH, F, Cl and N_3_) at C14 position through hydrofunctionalization. Furthermore, the synthetic utility of this powerful chemoenzymatic methodology was demonstrated by performing a 7-step semisynthesis of periplogenin and the diversity-oriented synthesis of cardenolide (+)-digitoxigenin and its three diastereomers in a concise manner.

---

Steroids are the second most marketed pharmaceuticals after antibiotics.^1^ C14-functionalized steroids, including C14α-OH and C14β-OH steroids, comprise a class of steroids of important bioactivities.^2-8^ Several members of this family have been utilized as medicines. For instance, proligestone is for animals’ estrus control,^5^ digoxin has been clinically used as a cardiotonic drug for treating heart failure through its positive inotropic effect^6^ (**Fig. 1a**) and so on (**Supplementary Fig. 1**). C14-OH steroids are widely distributed in nature, such as cardenolides and bufadienolides, the two major classes of C14β-OH steroids (>200 members) produced mainly by plants and toads respectively, albeit in very low yields.^2-4^ Besides the naturally occurring C14-OH steroids, various bioactive C14-functionalized steroids also have been synthesized for new drug discovery purpose.^2-4,7^

**Fig. 1.**
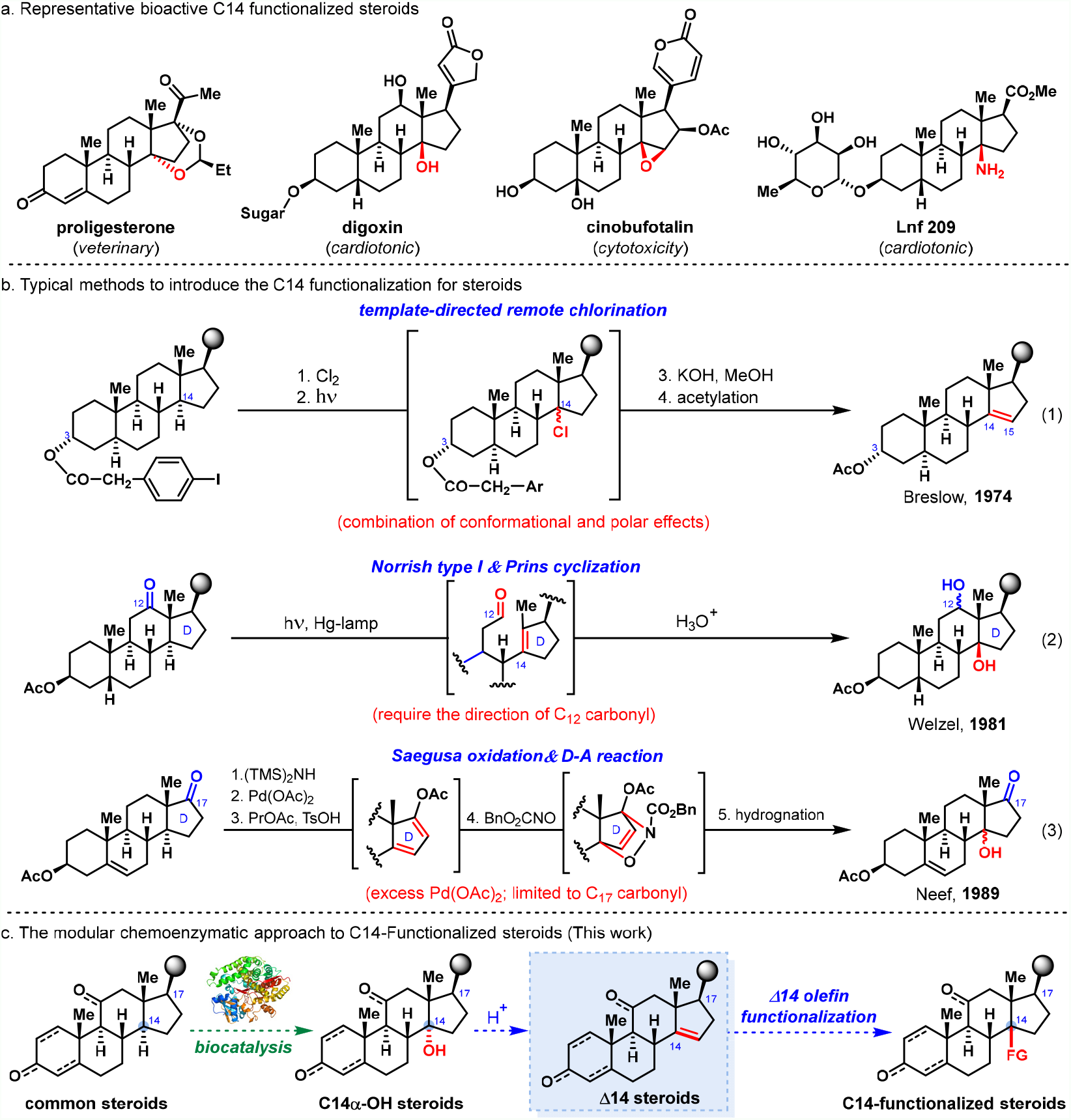
Approaches to access C14-Functionalized steroids.

Due to their significance for pharmaceutical utility, but scarcity from natural sources, syntheses of C14-functionalized steroids have become an area of sustained interest in organic chemistry.^8^ However, the inert reactivity of the tertiary C14−H bonds of steroids and the associated steric hindrance make their chemical functionalization a very challenging task. Substantial efforts have been devoted to addressing this issue in the past decades, culminating in the development of several impactful synthetic strategies,^4,9-15^ including (a) the elegant template-directed remote C14-H chlorination and elimination protocol to introduce Δ14 olefin by Breslow and coworkers (equation 1, **Fig. 1b**),^11^ (b) Norrish-type I/Prins cyclization protocol to introduce C14β-OH for 12-oxo-steroids by Welzel and coworkers (equation 2, **Fig. 1b**),^12^ (c) Saegusa oxidation & hetero-D-A reaction protocol to introduce C14-OH by Neef and coworkers (equation 3, **Fig. 1b**),^13^ (d) Saegusa oxidation and Mukaiyama hydration protocol to introduce C14β-OH by Baran and coworkers,^14^ and (e) allylic oxidation.^15^ Despite their effectiveness, these methods suffer from certain limitations, including the requirement of particular conformation/polarity (a) or pre-functionalization at specific positions of the substrates (e.g. C12 or C17 carbonyl group, Δ7 or Δ15 olefin) (b, c, d, e), strict reaction conditions (a, b) and multi-step chemical transformations (c, d). In addition to the chemical methods, biocatalytic hydroxylation is a promising alternative to access C14α-OH steroids directly. A few filamentous fungi and two P450 hydroxylases have been identified with steroidal C14–H α-hydroxylation activity.^16-23^ These biocatalysts, especially the P450 enzymes overexpressed in *Saccharomyces cerevisiae* are very attractive in the preparation of C14α-OH steroids due to the easy operation, low cost and high step-economy features. However, their poor regiospecificity and low catalytic reactivity for the steroids with a C17-side chain,^21-23^ constitute a significant obstacle to their broad utility. Thus, the aforementioned limitations of the state-of-the-art methods severely impede their applications in preparing C14-functionalized steroids and the downstream biological and pharmaceutical studies. Currently, the accessible C14-substitution of steroids is mainly limited to OH, while other types of C14-substitutions are rare^2,3,7^ (**Fig. 1a**). Therefore, a general strategy to access diversified C14-functionalized steroids in a concise and divergent fashion is highly desirable.

Recently, chemoenzymatic synthesis has emerged as an attractive strategy in the synthesis of complex natural products and medicines.^24-29^ It harnesses the power of both biocatalysis and chemical synthesis, thus can enormously shorten the synthetic routes and increase the overall efficiency. Inspired by the elegant chemoenzymatic studies,^30-33^ we envision a similar strategy to prepare the C14-functionalized steroids, which involves the identification of a versatile C14α-hydroxylase to prepare diversified C14α-OH steroids^34^ with a C17 substitution directly and the following chemical manipulations of the C14α-OH group to access other C14-functionalized steroids (**Fig. 1c**). We reason the C14α-OH can be readily transformed into Δ14 olefin through elimination, and the newly formed Δ14 olefin can serve as a versatile handle to install diversified functional groups at C14 position through its hydrofunctionalization.^35^ From a medicinal chemistry perspective, this chemoenzymatic strategy will be able to provide an ideal synthetic blueprint to target a wide spectrum of C14-functionalized steroids from common steroidal materials. Nevertheless, multiple challenges associated with the realization of this strategy are foreseeable. For instance, although the C17 substitution is pivotal for most of the bioactive steroids and related medicines, currently there are lack of efficient C14α-hydroxylases for these C17-substituted steroidal substrates.^16-23^ In addition, at the Δ14 olefin hydrofunctionalization stage, a complicated chemoselectivity issue may arise from the reactive functional groups of the A/B/C rings and side chain.^35^

Herein, we describe a modular chemoenzymatic approach to access C14-functionalized steroids. A novel C14α-hydroxylase (CYP14A) from *Cochliobolus lunatus* is identified. The engineered form of this enzyme is highly regioselective, efficient and permissive, and able to synthesize a variety of C14α-OH steroids with a variable C17 side chain. By combining with downstream chemical transformations, the C14α-hydroxyl group can be readily transformed into Δ14 olefin and other important functional groups (e.g. epoxide, β-OH, F, Cl and N_3_). The power of this strategy is demonstrated by a 7-step semisynthesis of periplogenin and diversity-oriented synthesis of cardenolide (+)-digitoxigenin and its three diastereomers in a concise manner.

## Results

### Identification of the regiospecific C14α-hydroxylase

The reported C14α-hydroxylase (P450_lun_) from *Cochliobolus lunatus* ATCC™ 12017 is poorly specific toward C17-substituted steroids, which converts cortexolone (also names as RSS) into a mixture of C14α- and C11β-hydroxylation products (14α-OH-RSS: 11β-OH-RSS = 2:3).^22^ To discover the hydroxylase with improved regiospecificity, we first explored the C14-hydroxylation activity of the three laboratory housed *C. lunatus* strains (CGMCC 3.4381, CGMCC 3.3589, JTU 2.406) using two common C17-substituted steroids progesterone (**1a**) and deoxycorticosterone (**1b**) as the substrates. Although CGMCC 3.4381 produced a mixture of C14α- and C11β- as well as other uncharacterized hydroxylated products (1:1 and 3:2 for **2a/2a′** and **2b/2b′** respectively, **Fig. 2a** trace I and IV), the rest two strains exhibited an improved regiospecificity, particularly for CGMCC 3.3589 which can specifically produce the C14α-hydroxylated products **2a** and **2b** only except for a minor dihydroxylated derivative (**2a′′**) of **1a**. (**Fig. 2a**, trace II and V). We next cloned the C14α-hydroxylase genes from the three strains (see **Supplementary methods**). The hydroxylase from CGMCC 3.4381 is identical to the reported P450_lun_, while the C14α-hydroxylases (CYP14A) from CGMCC 3.3589 and JTU 2.406 are the same which shows 82 % identity to P450_lun_.

**Fig. 2.**
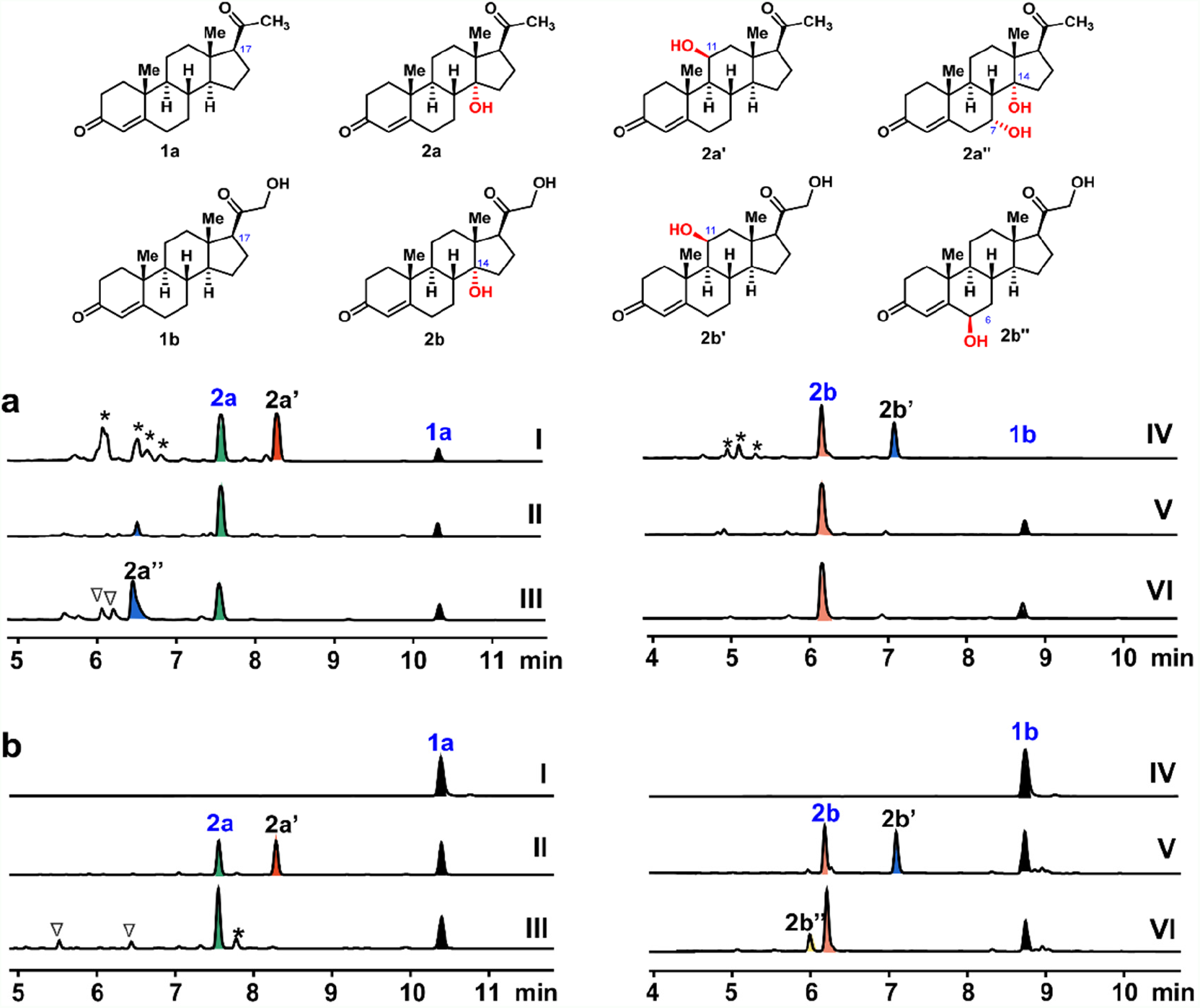
UPLC analysis of the C14α-OH steroids produced by biocatalysis. **a**, biocatalytic hydroxylation of steroids (100 mg L^-1^) by *C. lunatus* in the fermentation media (pH 6.1) for 72 hrs. I) CGMCC 3.4381, fed with **1a**; II) CGMCC 3.3589, fed with **1a**; III) JTU 2.406, fed with **1a**. IV) CGMCC 3.4381, fed with **1b**; V) CGMCC 3.3589, fed with **1b**; VI) JTU.2.406, fed with **1b**; Peaks with asterisk and triangle are uncharacterized monohydroxylated and dihydroxylated products respectively. **b**, biocatalytic hydroxylation of steroids (100 mg L^-1^) by *S. cerevisiae* in 50 mM PBS buffer (pH 7.2) for 3 hrs. I) *S. cerevisiae* YPH499 harboring empty vector, fed with **1a** (control); II) *S. cerevisiae-* P450_lun_, fed with **1a**; III) *S. cerevisiae*-CYP14A, fed with **1a**. IV) *S. cerevisiae* YPH499 harboring empty vector, fed with **1b** (control); V) *S. cerevisiae-*P450_lun_, fed with **1b**; VI) *S. cerevisiae*-CYP14A, fed with **1b**.

These two genes were further codon-optimized and overexpressed in the *Saccharomyces cerevisiae* YPH499. The resulting strains were utilized as a whole-cell biocatalyst to evaluate the hydroxylation activity of **1a** and **1b** (**Fig. 2b**). Like CGMCC 3.4381, *S. cerevisiae*-P450_lun_ was poorly regiospecific toward both substrates and produced mixtures of **2a/2a′** (1:1) and **2b/2b′** (5:4) (**Fig. 2b**, trace II and V). This performance was also similar to the biotransformation result of RSS,^22^ suggesting that its poor regiospecificity toward C17 substituted steroids was a general characteristic. *S. cerevisiae*-CYP14A exhibited a slightly different specificity from its fungal systems (**Fig. 2b**). The recombinant strain produced a few minor hydroxylated/dihydroxylated byproducts of **1a** (23% in total) and a 6β-OH byproduct of **1b** (**2b′′**) (27%), but not the byproducts of the fungal system. Co-expression of the *C. lunatus* P450 reductase (from *C. lunatus* ATCC™ 12017^22^) with CYP14A showed a similar performance (**Supplementary Fig. 2**). These clues suggested that **2a′′** and the related byproducts in CGMCC 3.3589/ JTU 2.406 were not produced by CYP14A. At the same time, overexpression of CYP14A in *S. cerevisiae* lead to a slight decrease in specificity since the new byproducts were not detected in the control strain (**Fig. 2b**). We speculated that the slight decrease in regiospecificity might be due to the high-level expression of CYP14A, which magnifies the side reactivities. Overall, despite the slight decrease in catalytic specificity, *S. cerevisiae*-CYP14A was still highly regiospecific and much superior to P450_lun_ in the synthesis of **2a** and **2b**. This yeast system, reducing the conversion time by 7 to 8 times, was far more efficient than the original fungal system, which provided a very effective and robust platform for steroids C14α-hydroxylation.

### Engineering CYP14A to improve its regiospecificity and catalytic reactivity

We next evaluated the substrate specificity of CYP14A by assaying a series of representative C17-substituted steroidal substrates. Except for the uncommon α-configuration of C17 and β-configuration of C5, CYP14A (in *S. cerevisiae*) can well tolerate the variations in the C17 side groups and oxidation state in A ring (**Table 1**). It can transform the substrates **1a–f** into the corresponding C14α-hydroxylated products in a medium to good regiospecificity (57−92%) and conversion (45−70%). The broad specificity made it desirable to synthesize C14α-OH steroids in a direct manner efficiently. To identify the critical residues for regiospecificity, the protein structure of CYP14A was built using RobeTTaFold^36-37^ (see **Supplementary Fig. 3**), and then all the 18 residues in the binding cavity were mutated into alanine and assayed for the C14α-hydroxylation activity of **1b**. Gratifyingly, four mutations remarkably alter the regiospecificity, among which I111A and V124A improved the C14-specificity from 73.4% (by WT) to 84.7% and 80.1% respectively, while M115A and F297A significantly decreased the C14-specificity (**Table 1** and **Supplementary Fig. 4**). Other mutations either abolished catalytic activity (H301A, T302A, F368A, K489A) or showed no apparent change to the ratio of **2b′′**/**2b** (F107A, P108A, R121A, T123, V213A, M305A, P363A, V364A, S367A, T369A and V490A) (**Supplementary Fig. 4**).

**Table 1.** Biocatalytic synthesis of C14α-OH steroids.

| Enz. | WT |  | I111A |  | V124A |  | I111A-V124A |  | M115K |  |
| --- | --- | --- | --- | --- | --- | --- | --- | --- | --- | --- |
| Sub. | C (%)<sup>a</sup> | S (%)<sup>b</sup> | C (%) | S (%) | C (%) | S (%) | C (%) | S (%) | C (%) | S (%) |
| **1a** | 48.5 | 77.3 | **62.1** | **83.7** | 49.3 | 76.4 | 54.3 | 78.2 | 41.2 | 63.7 |
| **1b** | 65.3 | 73.4 | 71.4 | 84.7 | 70.4 | 80.1 | 70.3 | 85.2 | **73.7** | **88.2** |
| **1c** | 61.5 | 85.2 | 67.7 | 88.4 | 65.5 | 86.3 | 65.4 | 86.2 | **68.4** | **90.5** |
| **1d** | 56.2 | 69.0 | 37.4 | 45.1 | 52.5 | 70.8 | 21.6 | 54.4 | **70.5** | **85.3** |
| **1e** | 45.3 | 57.4 | 15.2 | 20.5 | 44.7 | 58.4 | 11.3 | 31.6 | **78.5** | **92.2** |
| **1f** | 70.3 | 92.1 | 65.5 | 90.3 | **74.6** | **94.2** | 59.2 | 85.3 | 64.3 | 86.5 |

<sup>a</sup>C indicates HPLC yield of C14α-OH products **2a–f**; <sup>b</sup>S indicates C14α-hydroxylation specificity.

Docking **1b** into the binding cavity of CYP14A (**Fig. 3**) revealed that I111 and V124 are at the bottom of the binding pocket. Their mutation into small residue can enlarge the binding cavity and let the steroidal framework slightly go deeper to the pocket to facilitate the approach of C14α-hydrogen to the iron of heme (**Fig. 3b**). With this clue, other large-to-small mutations including I111G, I111V, V124G and I111A-V124A were constructed and evaluated, among which I111A-V124A showed a further improvement in C14-hydroxylation specificity (85.2%) (**Table 1** and **Supplementary Fig. 6**).

**Fig. 3.**
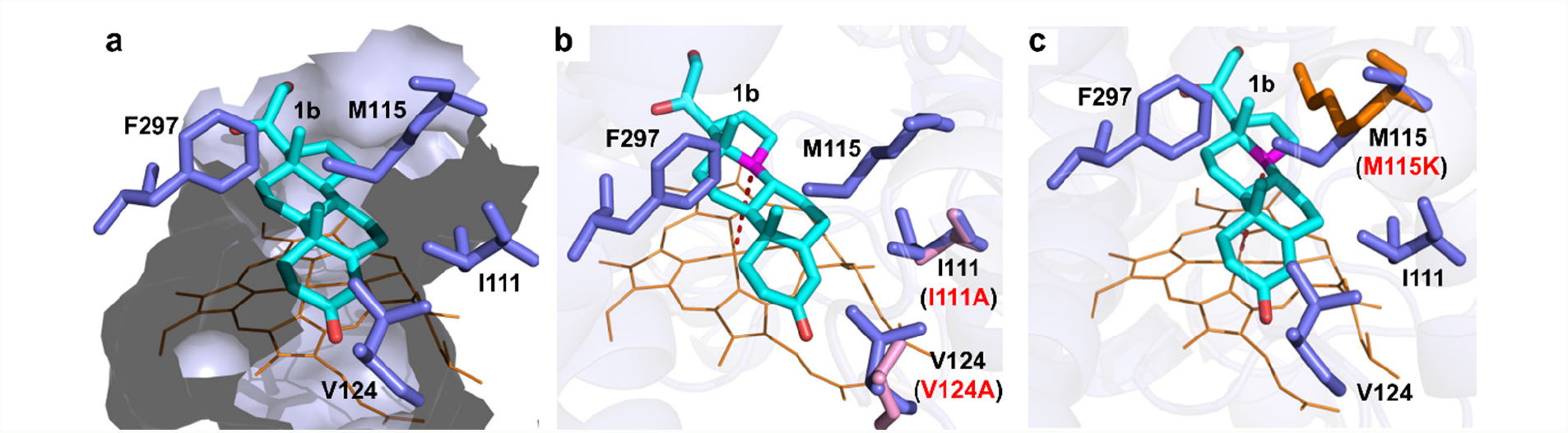
In silico model of CYP14A with the substrate 1b. **a**, the binding cavity of CYP14A. The substrate **1b**, heme, and the binding cavity are colored in cyan, brown and light blue, respectively. The four critical residues (in blue) are labeled by the corresponding amino acids. **b**, comparison of the wild-type enzyme and I111A-V124A. The mutated alanine residues and the C14 of the substrate are colored in pink. **c**, comparison of the wild type enzyme and M115K. The mutated lysine residue and the C14 of the substrate are colored by brown and pink, respectively.

M115 and F297 residing at the top of the substrate were essential for maintaining the C14-hydroxylation conformation by pushing the steroidal framework of **1b** to the iron of heme (**Fig. 3**). To explore the potential of other residues, they were mutated into either large or small residues. Although most mutants were similar or inferior to the wide type enzyme, M115K remarkably improved the C14α-selectivity/conversion from 65.3/73.4 to 73.7%/88.2% (**Table 1, Supplementary Fig. 5**), assumably via pushing the steroidal framework closer to the heme (**Fig. 3c**). Lastly, the beneficial mutations were combined, while their simultaneous mutation was either neutralizable or detrimental to the specificity or activity, leading to decreased regiospecificity (M115K-V124A) or abolished catalytic activity (M115K-I111A and M115K-I111A-V124A) (**Supplementary Fig. 6**). Together, these findings successfully improved the regiospecificity and catalytic activity of CYP14A and gave a clear clue to further rational engineering of CYP14A to obtain the more efficient and versatile biocatalysts.

With the specificity-improved mutants, we further applied them to transform other substrates (**1a, 1c-f**). To our delight, these four mutants all demonstrated improved regiospecificity and conversion to most substrates (**Table 1, Supplementary Fig. 7**). Among them, I111A, M115K and V124A were the most effective ones in improving the catalytic specificity for **1a, 1b**-**1e** and **1f**, respectively; and most remarkably, the specificity and conversion of the highly useful steroidal precursor **1e** were improved by 35% and 32% respectively by M115K comparing to the wild-type enzyme. With the best mutants, each substrate’s the catalytic specificity and conversion can reach 83−94% and 62.1-78.5% respectively by a 6-12 hrs biotransformation. Taken together, these engineered P450 hydroxylases were far better than the previously reported biocatalysts^21-23^ in regiospecificity, substrate promiscuity and catalytic reactivity, which enabled a highly efficient and practical biocatalytic way to the synthesis of C14α-hydroxylated steroids with a C17 substitution.

### Preparing Δ14 olefinated steroids from C14α-OH steroids

Since the enzymatic C14−H α-hydroxylation has become a reliable and efficient method to assemble the C14-α-OH steroids, a following chemical exploration could be carried out to establish the modular platform for obtaining novel C14−functionalized steroids. Initial attempts on direct replacement of C14-α-OH group of **2a**–**f** to install new C14 functionality were unsuccessful. Instead, dehydration of the C14 tertiary alcohol to form Δ8 or Δ14 olefin under acidic (or basic) conditions was the major reaction pathways. In addition, the C17 side chains of **2a**–**f** were also susceptible to multiple competing side reactions as well. Nevertheless, we surmised if the Δ14 olefin product can be obtained from the C14-α-OH intermediate chemoselectively, the newly formed olefin motif could be utilized as a versatile transformative handle for late-stage functionalization at the C14 position. Thus, we selected C14-hydroxy progesterone **2a** as the model substrate to investigate the dehydration reaction. Gratifyingly, after extensive reaction conditions optimization (see Supplementary **Table 1** for details), trifluoroacetic acid (TFA) was identified to be the optimal promoter for dehydration to afford the desired **3a** in the highest yield (95%).

To our delight, the identified optimal dehydration conditions can be applied to other C14-α-OH steroids in **Table 1** to prepare the corresponding Δ14 olefinated steroids (**Scheme 1A**). For instance, under the standard dehydration conditions, **2b**−**d** were transformed into olefins **3b**−**d** in good to excellent yields (67−99%). However, only a moderate yield (42%) was obtained for **2e**, while no elimination took place for **2f**. It was probably because TFA was not acidic enough to promote the desired dehydration in these two cases. Indeed, when **2e** was treated with a stronger Brønsted acid *p*-toluenesulfonic acid (TsOH•H_2_O) under microwave irradiation, **3e** was formed in a much higher yield (68%, 90% brsm). In contrast, dehydration of **2f** was facilitated by the Lewis acid BF_3_•Et_2_O to afford the dehydrated product **3f** as a separable mixture of C17 epimers (68% combined yield, β/α = 2:1).

### Synthesis of C14 functionalized steroids

Owing to the versatile reactivity of the olefin group,^35,38^ we then extended our efforts to the installation of diversified C14 functional groups through the elaboration of the obtained Δ14 olefinated steroids **3a**−**f**, which was demonstrated in **Scheme 1B**. At first, the model substrate **3a** was stereospecifically oxidized by *m-*CPBA to give the α-configurated epoxide **4aa** in almost quantitative yield. Interestingly, the stereochemistry of epoxidation is completely reversed for **3a** as compared to the C17 carbonyl counterpart.^9^ Next, subjecting **3a** to Fe(acac)_3_-catalyzed anaerobic Mukaiyama conditions using nitrostyrene as the external oxidant^39^ afforded the hydrated product **4ab** as a mixture of C14 diastereomers in almost quantitative yield (β/α = 1.3:1). In addition, the intriguing C14 azide group can be introduced through a Fe^III^/NaBH_4_-mediated free radical hydroazidation^40^ of **3a** to deliver **4ac** as a mixture of C14 diastereomers in 78% yield (β/α = 1:1.8). Similarly, using Selectfluor as the fluoro atom transfer reagent, a Fe^III^/NaBH_4_-mediated free radical hydrofluorination^41^ was realized to afford **4ad** as a mixture of C14 diastereomers in 72% yield (β/α = 1:1). Moreover, by utilizing the *in situ* generated HCl as the reagent, a classic electrophilic hydrochlorination of **3a** was achieved with excellent regioselectivity and diastereoselectivity, and only Markovnikov addition product **4ae** was isolated in 60% yield as a single diastereomer with the β-configuration. Interestingly, the configuration of C17 stereocenter was completely epimerized in this transformation, as revealed by X-ray crystallographic analysis of **4ae** (CCDC 2116722). In order to demonstrate the reliability and robustness of the olefin derivatization strategy, we also tested the olefinated steroid **3c** with a different A/B ring conformation compared to **3a**. Gratifyingly, the same transformations were successfully carried out on **3c** to afford the corresponding products with comparable results regarding reaction efficiency and diastereoselectivity. Notably, the C3 ketone moiety of **3c** was more prone to be reduced than **3a** under the hydroazidation conditions, thus the corresponding product **4cc** was obtained in the reduced form with a C3-α-OH configuration. Additionally, radical hydrofluorination of **3c** delivered **4cd** as a single diastereomer with the 14-β-configuration in 60% yield. It is worth mentioning that all the formed C14 diastereometic products (**4ab**−**ad** and **4cb**− **cc**) were readily separated by flash column chromatography and characterized independently. The absolute configurations of **2b, 4aa, 4cc** and **4ae** were unambiguously confirmed by X-ray crystallographic analysis (Scheme 1B), and those of other products were assigned by analogy after careful NMR analysis. Therefore, the olefin derivatization strategy for the installation of C14 functionality was proven to be very successful, particularly in preparing novel C14 functionalized steroid analogs not accessible by other alternative methods.^9,10^ We believe these intriguing C14 functionalized steroid analogs will be very useful in complex steroid synthesis and new drug discovery. For instance, the C14-β-OH diastereomers of **4ab** and **4cb** can serve as useful common intermediates for the synthesis a series of important natural C14-β-OH steroids.^4,9,10^

**Scheme 1.**
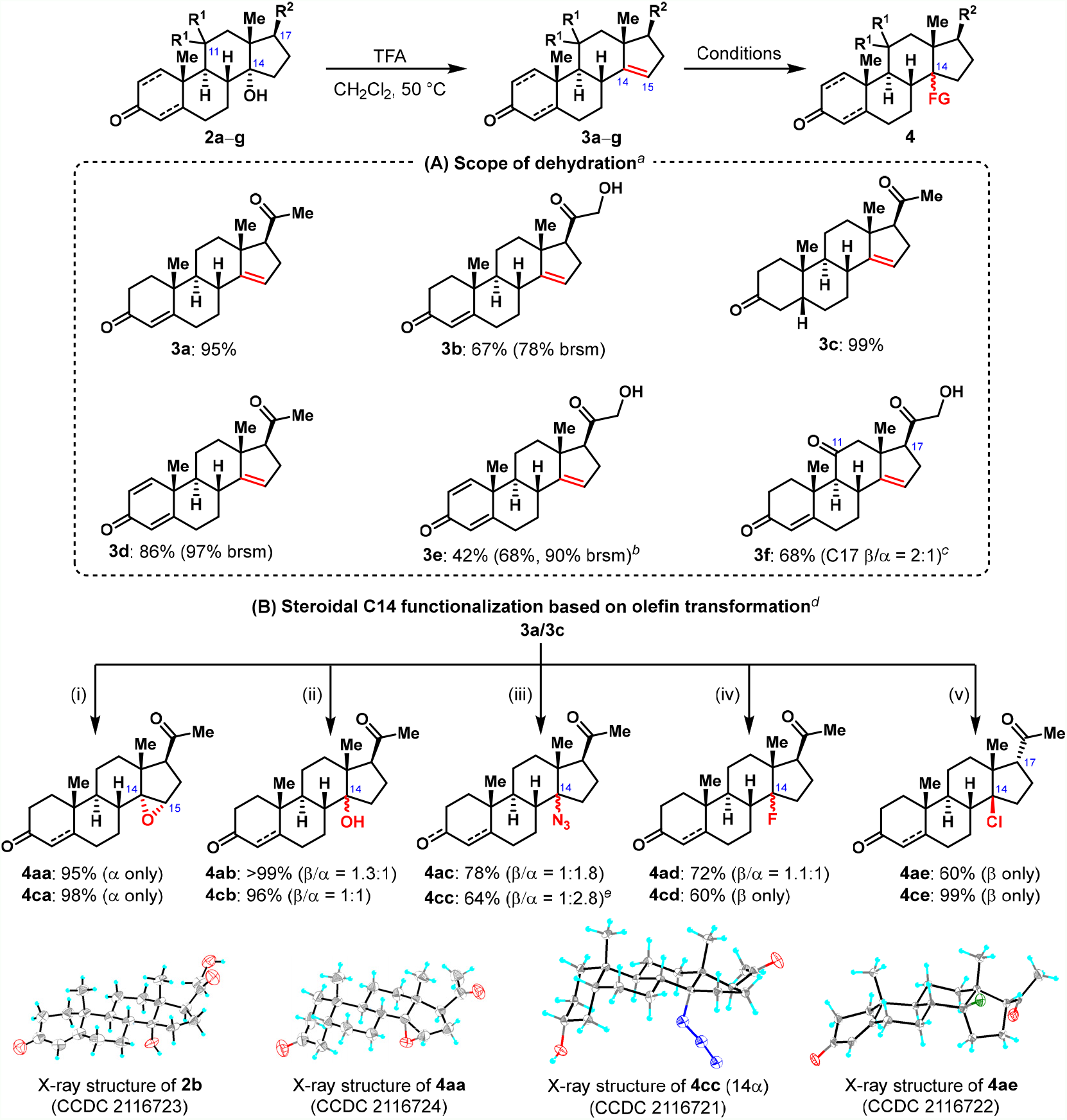
Dehydration of C14α hydroxy and olefin derivatization. ^*a*^Reaction was performed on a 0.03 mmol scale, and isolated yield was reported. ^*b*^Conditions: TsOH•H_2_O (3.0 equiv), H_2_O, microwave, 140 °C. ^c^Conditions: BF_3_•Et_2_O (1.0 equiv), CH_2_Cl_2_, 0 oC. ^*d*^Reaction was carried out on a 0.06 mmol scale. Isomers were isolated by flash column chromatography. ^*e*^C3 ketone of **3b** was reduced to C3 α-OH. Reaction conditions: (i) *m*-CPBA (1.2 equiv), CHCl_3_, r.t. (ii) Fe(acac)_3_ (10 mol%), PhSiH_3_ (4.0 equiv) nitrostyrene (2.0 equiv), EtOH, 60 °C. (iii) Fe_2_(ox)_3_•6H_2_O (4.0 equiv), NaBH_4_ (10.0 equiv), NaN_3_ (3.0 equiv), EtOH, H_2_O, 0 °C. (iv) Fe_2_(ox)_3_•6H_2_O (4.0 equiv), Selectflour (4.0 equiv), NaBH_4_ (10.0 equiv), MeCN/THF/H_2_O, 0 °C to r.t.. (v) AcCl, EtOH, r.t..

### Semisynthesis of periplogenin

To demonstrate the synthetic utility of the powerful methodology, a concise semi-synthesis of complex natural product periplogenin from intermediate **3b** was performed (**Fig. 4**). Since the α-hydroxyl ketone side chain of **3b** was rather reactive, it was first transformed into the relatively stable butenolide motif through reaction with Bestmann ylide,^42^ affording **5** in 90% yield. The following Mn(acac)_2_/PhSiH_3_-mediated hydration^43,44^ of **5** delivered the desired 14β-hydroxy intermediate **7** in 43% yield together with an equal amount of the C14α-epimer **6**, which can be recycled through BF_3_•Et_2_O mediated dehydration. The following synthetic challenge is the introduction of a 5β-hydroxy group. Initial attempts on the direct epoxidation of **7** proved unsuccessful due to the interference of the butenolide motif and lack of good diastereocontrol. Therefore, we chose the following indirect epoxidation strategy. The diastereoselective reduction of C3 carbonyl of **7** by NaBH_4_ led to C3-β-OH **8** in 82% yield, which was then reacted with *m-*CPBA to deliver the β-epoxidated product **9** exclusively. The following Dess-Martin oxidation and regiospecific cleavage of the epoxide^45^ protocol afforded **11** in good yield. Finally, a diastereoselective reduction of C3 carbonyl of **11** by L-selectride delivered periplogenin (**12**) in 71% yield. Thus, the protecting-group-free synthesis^46^of periplogenin was achieved in just seven steps from **3b**.

**Fig. 4.**
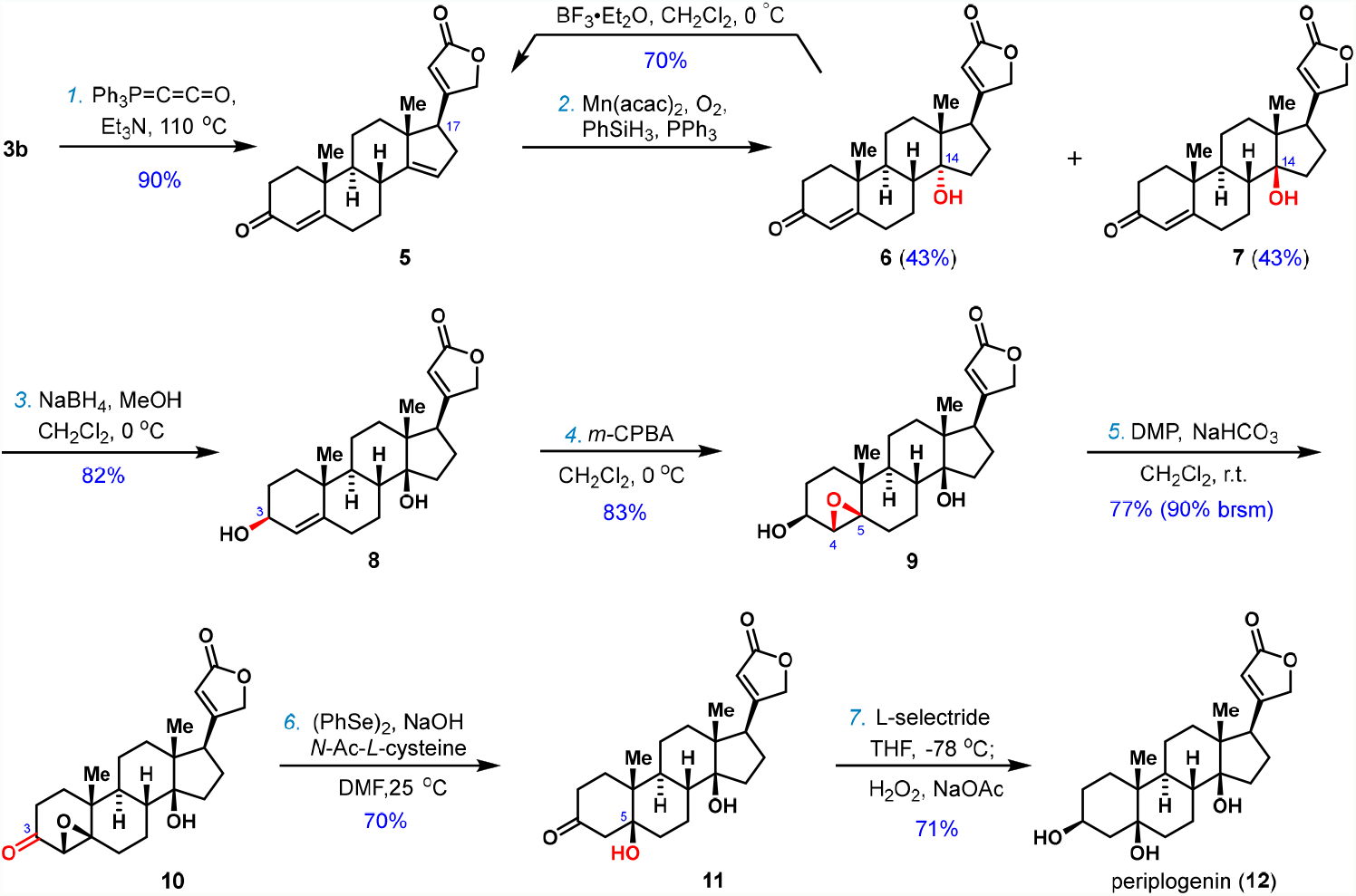
Facile synthesis of periplogenin.

### Synthesis of (+)-digitoxigenin and its three diastereomers

Lastly, to demonstrate the value of our chemoenzymatic strategy in medicinal chemistry, a diversity-oriented synthesis^47,48^ of cardenolide (+)-digitoxigenin and its three diastereomers was performed (**Fig. 5**). (+)-Digitoxigenin is the aglycon of digitoxin as well as the synthetic precursor of cardiotonic drug digoxin. (+)-Digitoxigenin itself exhibits antileishmanial (IC_50_ = 18.4 μM) and anticancer activities.^49^ The synthesis commenced with **2b**, which was prepared from **1b** on a gram scale under the optimal enzymatic conditions by *S. cerevisiae*-M115K (**Table 1**). At first, **2b** was subjected to Pd/C-catalyzed hydrogenation to solely afford A/B *cis*-fused intermediate **13** in 86% yield. Following TFA-promoted dehydration took place smoothly to deliver olefin **14**, which was readily transformed into the critical butenolide intermediate **15** after treatment with Bestmann ylide. Subsequent reduction of C3 carbonyl of **15** with L-selectride at −78 °C produced the C3-β-OH configurated **16** in 82% yield. Ultimately, a Mn(acac)_2_/PhSiH_3_-mediated hydration of **16** afforded (+)-digitoxigenin (**17**) and its C14 epimer 14α-hydroxydigitoxigenin (**18**) as a 1:1 separable mixture in 84% combined yield. Interestingly, the reduction of C3 carbonyl of **15** by NaBH_4_ led to C3-α-OH **19** as the major product (65% yield). A subsequent Fe(acac)_3_-catalyzed hydration afforded 3α-dihydroxyldigitoxigenin (**20**) and 3α,14α-dihydroxyldigitoxigenin (**21**) as a 1:1 separable mixture in 70% combined yield. Thus, the protecting-group-free syntheses of (+)-digitoxigenin and its three diastereomers were achieved in just six steps from **1b**. Moreover, as revealed in **Scheme 1B**, other analogs of digitoxigenin can be accessed divergently from intermediates **16** and **19** through late-stage hydrofunctionalization of Δ14 olefin. We envision that this synthetic route’s concise and divergent features will make it attractive for developing new steroidal medicines via careful SAR studies of these accessible steroidal analogs.

**Fig. 5.**
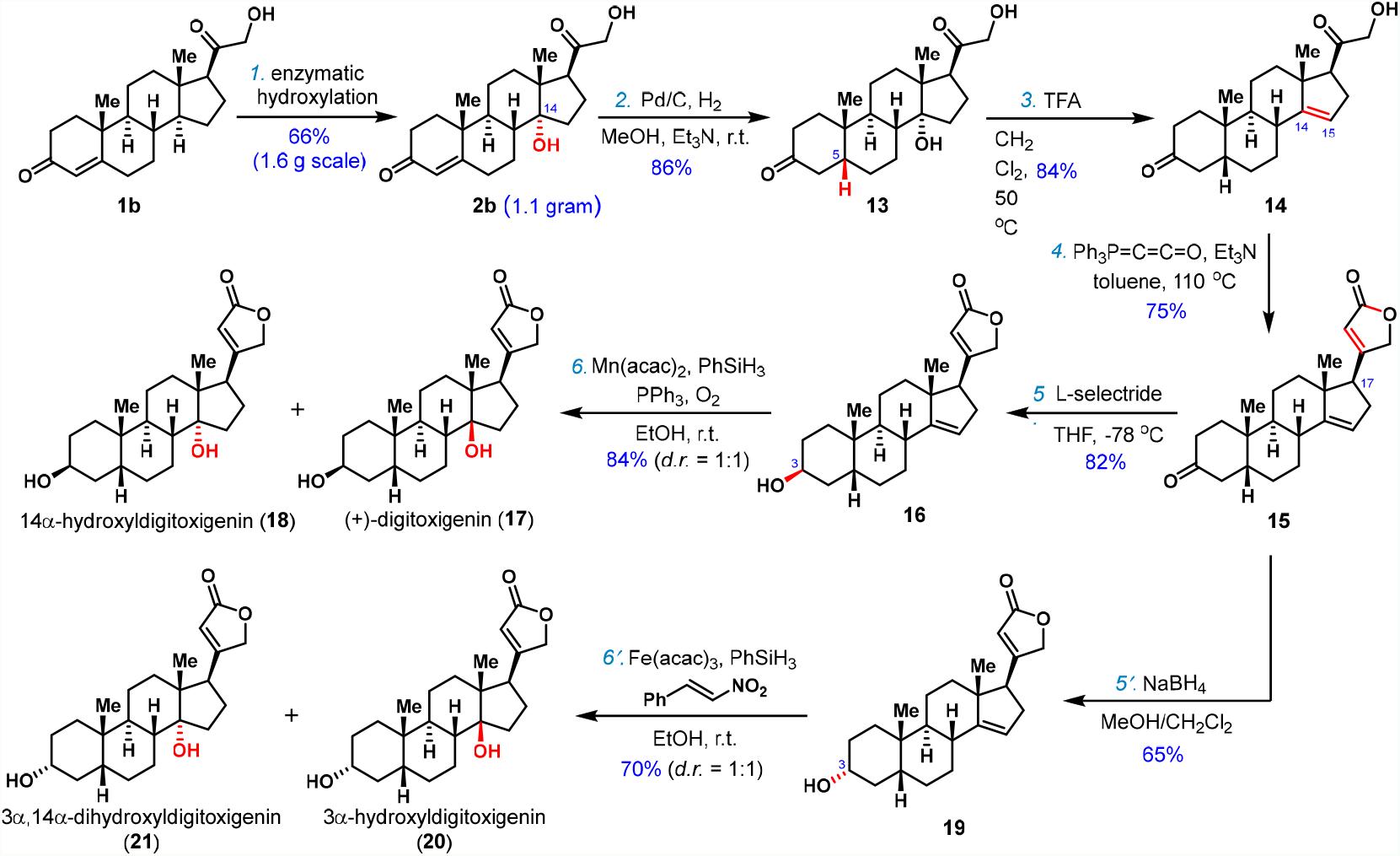
Diversity-oriented synthesis of (+)-digitoxigenin and its three diastereomers.

## Conclusions

In summary, we have developed a modular and efficient chemoenzymatic approach to access diversified C14-functionalized steroids. First, a novel C14α-hydroxylase (CYP14A) from *Cochliobolus lunatus* is identified. The engineered form of this enzyme is highly regioselective, efficient and permissive. Based on this reliable and efficient biocatalytic method, a range of C14α-OH steroids with C17 side chain are prepared in good yields, which are then transformed into Δ14 olefin through a facile elimination. The newly formed Δ14 olefin serves as a versatile handle to install diversified functional groups (e.g., epoxide, β-OH, F, Cl and N_3_) at C14 position through hydrofunctionalization. Furthermore, the synthetic utility of this powerful chemoenzymatic methodology is demonstrated by performing a 7-step semisynthesis of periplogenin and the diversity-oriented synthesis of cardenolide (+)-digitoxigenin and its three diastereomers in a concise manner. We believe this chemoenzymatic approach will provide a robust platform for the subsequent medicinal chemistry exploration of new steroidal drugs.

## Data availability

Crystallographic data for compounds **2b, 4aa, 4cc** and **4ae** are available free of charge from the Cambridge Crystallographic Data Centre (CCDC) under reference numbers 2116723, 2116724, 2116721, and 2116722, respectively. All other characterization data and detailed experimental procedures are available in the supplementary materials. The sequence of the P450 genes reported in this paper has been deposited in GenBank under the accession numbers: ON605617 (CYP14A from CGMCC 3.3589 and JTU2.406) and ON605618 (P450lun, from CGMCC 3.4381).

## Acknowledgements

We thank Dr. X.-G. Meng (CCNU) for X-ray crystallographic analysis assistance, Prof. L. Shao (Shanghai Institute of Pharmaceutical Industry) for kind gifting *C. lunatus* JTU 2.406, Prof. H.-G. Cheng, Mr. T. Yang, Mr. J.-X. Ye, Mr. L. Zhou and Ms H. Wei (WHU) for helpful discussions, Dr. L. Cao (WHU) for assistance with the preparation of the manuscript. This work was supported by the National Key R&D Program of China (2018YFA0900400, X.Q.), the Fundamental Research Funds for the Central Universities (2042021kf0214, Q.Z.) and the start-up funding from Wuhan University.

## Author contributions

Q.Z. and X.Q. conceived this project. F.S., M.Z., J.W., H. L., Z. L. and B. L. performed the experiment under the supervision of X.Q. and Q.Z.. F.S., M.Z., Q.Z. and X.Q. co-wrote the manuscript. All authors discussed the results and commented on the manuscript.

## Table of contents graphic

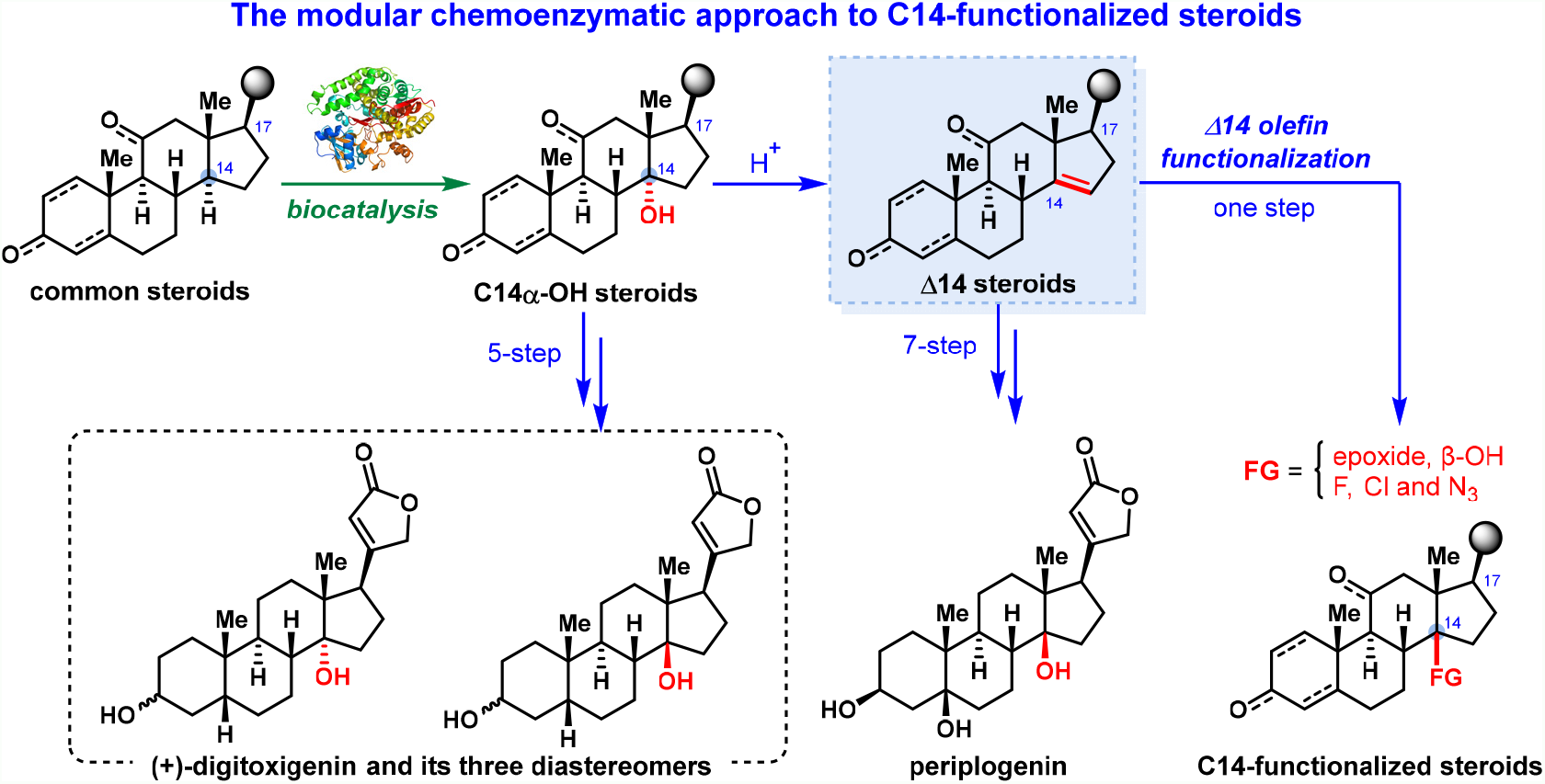

